# Transcriptome and Proteome analysis of *Hemidactylus frenatus* during initial stages of tail regeneration

**DOI:** 10.1101/2019.12.17.879718

**Authors:** Sai Pawan, Sarena Banu, Mohammed M Idris

## Abstract

Epimorphic regeneration of appendages is a complex and complete phenomenon found in selected animals. *Hemidactylus frenatus*, the common house gecko has the remarkable ability to regenerate the tail tissue upon autotomy involving epimorphic regeneration mechanism. This study has identified and evaluated the molecular changes at gene and protein level during the regeneration of tail tissue. Based on next generation transcriptomics and *De novo* analysis the transcriptome and proteome library of the gecko tail tissue was generated. A total of 417 genes and 128 proteins were found to be associated with the regeneration of gecko tail tissue upon amputation at 1, 2 and 5-day post amputation against control, 0dpa through differential analysis. The differentially expressed genes and proteins expressed a similar pattern for the commonly identified 36 genes/proteins involved in regeneration of the tail tissue. Similarly, the expression analysis of 50 genes were further validated involving real time PCR to authenticate the transcriptomics analysis. 327 genes/proteins identified from the study showed association for GP6 signaling pathway, Protein kinase A signaling, Telomerase signaling BAG2 signaling, paxiling signaling, VEGF signaling network pathways based on network pathway analysis. This study empanelled list of genes/proteins associated with tail tissue regeneration and its association for the mechanism.

## INTRODUCTION

Regeneration is a sequential process specifically controlled by cellular mechanisms to repair or replace tissue or organ from an injury. The mass of undifferentiated cells (blastema) surrounding the injured tissue results in the formation of fully functional replica which enacts the phenomenon of epimorphic regeneration [1]. The ability to regenerate a full limb is absent in mammals but urodeles; teleost and amniotes have the propensity to replace limbs, spinal cords, nervous system, heart, tail and, other body parts [2]. To understand the mechanistic framework of regeneration, researchers have been majorly associated with urodeles amphibians [3]; teleost fishes [4]. Though amniotes i.e. lizards are closely related to mammals and other vertebrates, very little attention has been given to carry out regeneration on this organism [2], [5].

Lizard capacity to self-detach or amputate their tail in flight response from predators is known as the shedding of the tail or caudal autotomy [6]. These stages include exudation of excessive blood loss; minimizing muscle and bone tissue damage; controlled vascularisation; promotes wound epithelium and further unsegmented remodelling [7]. The Detachment of tail activates multiple cellular responses which spurts the blastema mediated cell proliferation, angiogenesis, remodelling and generating a replacement. Studies related to the patterning of successful regenerated autotomized tail or replica consists mainly of nerve cells networking, unsegmented cartilaginous, muscle cells, blood vessels and differential remodelling of cells which make this an interesting research to comprehend the cellular and molecular mechanisms associated with the development [8]–[10]. With the elucidation of the genes and their associated proteins involved in regeneration, there can be a possibility of understanding why in humans regeneration is restricted as compared to amniotes. This also might pave in giving insights into spinal injuries and the fabrication of replacement therapies [11]. Research on the regeneration of tail and its governing molecular mechanism in other lizard species like Hemidactylus falviviridis [12]; Green Anole [13–14]; and other vertebrates axolotl [15–17] have been carried out related to histology, proteomics and genomics [12–17]. Though many genes or proteins have been known still the underlying mechanisms of these molecules remains ambiguous. The present study is novel on this species “*Hemidactylus frenatus*” [18] commonly known as “Common House Gecko”. They are widely accessible, have a short life span and can be housed with minimal requirements and maintenance conditions. These species have been studied generally for their behavioural, habitat diversification and regenerative abilities [7], [19–22].

Lately, molecular and gene-specific studies focus on the role of genes like SOX9, PAX7, BMP, FGF, Wnt, MMP’s which are usually associated with regeneration [23]. The expression of SOX9 during cartilage formation, PAX7 associating in tissue morphogenesis [24], BMP6 and other protein expressions were known [25, 26]. Constitutive expression and regulation of these genes have revealed a significant role during the developmental stages of regeneration. The slight alteration in the regulation of any of these genes results in a disrupted replacement. Thus, the activation and intricate cross-talk of these molecules during the process are still unaddressed and need to be better understood.

In this research, we sought to understand the molecular mechanisms of lost tail regeneration process during preliminary stages (0dpa, 1dpa, 2dpa, and 5dpa) which would further help in understanding the molecular cues and the genes or proteins associated with the process. The initial stages (i.e. wound healing) are responsible for wound epithelium and initiating apical epidermal cap (AEC) which regulates blastema formation [7]. Though, during the initial stages, the activity of the molecules regulating the regeneration process is not known which is crucial to instigate the replacement of the lost tissue. The study of prelude stages of regeneration has been carried out in zebrafish caudal fin [27]; other species [28] and shown changes in the regulation of genes or proteins. However, this stage-specific study has not been carried out in this species and as it is neccessary for deciphering the crucial molecules which kick starts the regenerative mechanism in these amniotes. Here, we list out the involvement of crucial genes or proteins and their differential expression associated with the molecular processes in tail regeneration through transcriptome and proteome approaches.

## MATERIALS AND METHODS

### 1. Animal and Sample Collection

The lizard, *Hemidactylus frenatus* were collected and maintained in a well-ventilated cage with adequate proper diet, optimum temperature (∼25°C) and the 12 hours light and dark cycle. Amputation of tail tissue were performed on batches of animals using sterile scalpel blade. Regenerating tail tissues preceding the amputation site were collected for each time points (0dpa, 1dpa, 2dpa, 5dpa) of (n=5). The collected control and regenerating tail tissues were washed with 1X phosphate-buffered saline (PBS), pooled and snap-frozen in liquid nitrogen and was store at −70°C until use. The animal experiment were performed in accordance with the protocol approved by the Institutional animal ethics committee of Centre for Cellular and Molecular Biology (IAEC/CCMB/Protocol # 66/2014).

### 2. Total RNA Isolation

Total RNA was extracted from the tissues of each time points using RNA isoPlus Reagent (Takara Bio, CA, USA) following the manufacturer’s protocol. The RNA yield and purity was calculated using NanoDrop 2000 Thermo fisher and gel analysis.

### 3. NGS transcriptomic analysis

The total RNA transcripts of each time point tissues were obtained based on next generation sequencing (NGS) analysis involving Illumina HiSeq 2000 [28]. All the transcripts obtained commonly from all the tissues were further assembled for De novo transcriptome analysis and functional annotation against non-redundant reptilian database using blastx [28]. Both the known and unknown gene sequences obtained from the NGS analysis were tabulated and submitted to NCBI and obtained accession number. Also the genes were translated for obtaining protein database.

### 4. Differential expression analysis

The transcripts were further analysed for their expression level using FPKM and analysed for differential expression level on 1, 2 and 5dpa time points against 0dpa as control [28]. Differentially expressed transcripts having at least 1.0 log fold change in any one of their regenerating time points were considered for the study.

### 5. Real Time PCR (RTPCR) analysis

Validation of fifty most significantly expressed genes having more than 2 log fold changes in any of its time points were selected for RTPCR analysis. Primers were designed using Primer3 software. Amplification of 14-3-3 protein zeta/delta isoform X1 and NADH dehydrogenase 1 alpha sub complex subunit 11 were used as housekeeping gene. RTPCR were performed in biological and technical replicates for each genes from the cDNA synthesized from 1 μgms total RNA using Takara SYBR green assay master mix. The relative expressions of the genes were estimated based on the RTPCR Ct value against control (0dpa) as the baseline.

### 6. Protein extraction and iTRAQ analysis

The total protein from the control and regenerating tail tissues were extracted using protein extraction buffer (7M urea, 2M thiourea, 18mM Tris-HCl, 4% CHAPS, 14mM Trizmabase, 2 Tablets EDTA protease Inhibitor, Triton X 0.2%, 50mM DTT) upon homogenization and sonication [27, 29]. The proteins were quantified using Amido black method [30] against BSA as standard. iTRAQ based quantitative proteomics analysis was performed in duplicates between the control (0dpa) and regenerating time points (1dpa, 2dpa and 5dpa) by loading 50 μg of total proteins in a 10% SDS-PAGE gel. The gel was stained, destained, documented and fractionated into four sequential groups. The gel fractions were washed; trypsin digested, labelled with isobaric tags iTRAQ 4-plex labelling and purified with the help of C-18 spin columns (Thermo Scientific). The purified/labelled peptides were vacuum dried and constituted in 5% acetonitrile (ACN) and 0.2% formic acid to the peptides for the LCMS/MSMS analysis [28]. The LCMS/MSMS run was performed in Orbitrap Velos Nano analyzer (Thermo) involving High Collision Dissociation (HCD) mode of acquisition with 50% normalized collision energy. The raw files were analyzed with Sequest HT proteome discoverer 1.4 (Thermo Scientific), with 1% FDR using percolator and XCorr (Score Vs Charge) against the NGS generated lizard database. All the proteins, in duplicates, were analyzed against the control (0dpa). Differentially regulated expression by more than 1-log fold changes were selected as proteins associated with regeneration.

### 7. Heat Map and Network Pathway analysis

All the differentially expressed genes and proteins were analysed for the heat map analysis involving heatmapper portal (www.heatmapper.ca) towards elucidating the hierarchical cluster analysis of the genes/proteins and the time points. The association of these genes / proteins were also analyzed for the association in network pathways based on Ingenuity pathway analysis (IPA).

## Results

### Transcriptomic analysis of regeneration

A total of 42551 transcripts sequences were obtained from the NGS analysis of all four tail tissue samples. All the sequences were obtained from a total of ∼ 30 million reads from each sample. De novo and functional annotation yielded 39953 transcripts with highest similarity with *Gekko japonicas* (41.3%) and no blast hit (49%) to any reptilian database. The transcripts were submitted to NCBI and obtained accession number (SUB6511228). The transcripts were further translated to obtain the protein sequence database for proteomic analysis.

A total of 417 genes were found to be associated with regeneration of gecko tail tissue for having at least one log fold change in one of its regenerating time points significantly. 254 genes were selected for further analysis such as heat map and network pathway analysis for having their expression pattern in all their regenerating time points (Figure 1a; Table 1). Genes such as Erythrocyte membrane protein (EPB41), retinoic acid receptor responder protein 1 and few uncharacterized proteins were found to be upregulated for having more than 5 log fold upregulation of gene expression. Similarly, Ryanodine receptor 1, Myosin 1B were found to be regulated austerely (Table 1). Clustal hierarchical heat map analysis of the differentially regulated genes revealed group of genes which were undergoing upregulation at 1dpa where found to be down regulated at 2 and 5dpa and vice versa with down regulated genes at 1dpa (Figure 1a). The analysis further reveals that 1dpa expression is out rooted with clustered 2 and 5dpa based on cluster analysis for the three regenerating time points against the control tissue expression.

**Figure 1:**
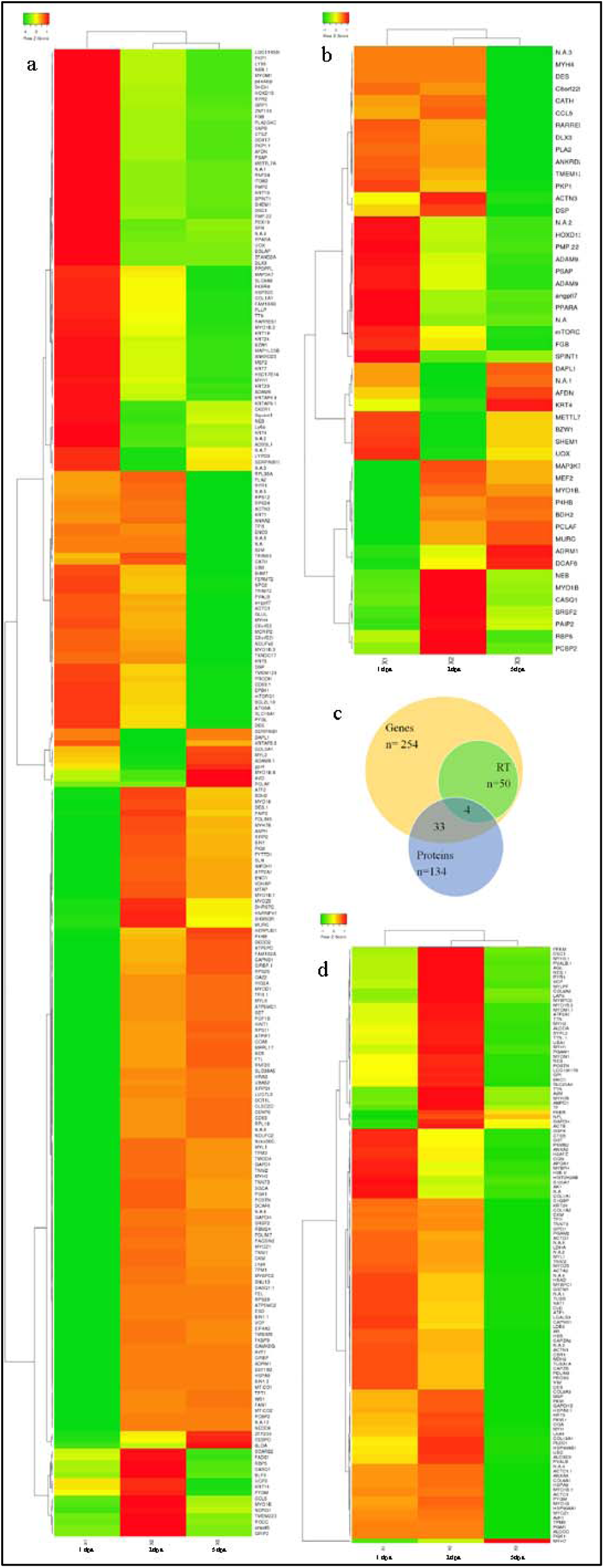
a. Heap map expression of *Hemidactylus frenatus* transcript during tail tissue regeneration b. Heat map expression of various genes differentially regulated during regeneration based on RTPCR analysis; c. Venn diagram for number genes identified from Transcriptomics, proteomics and Real time PCR based gene expression analysis D. Heat map expression of differentially expression proteins which are associated with tail regeneration.

**Table 1:**
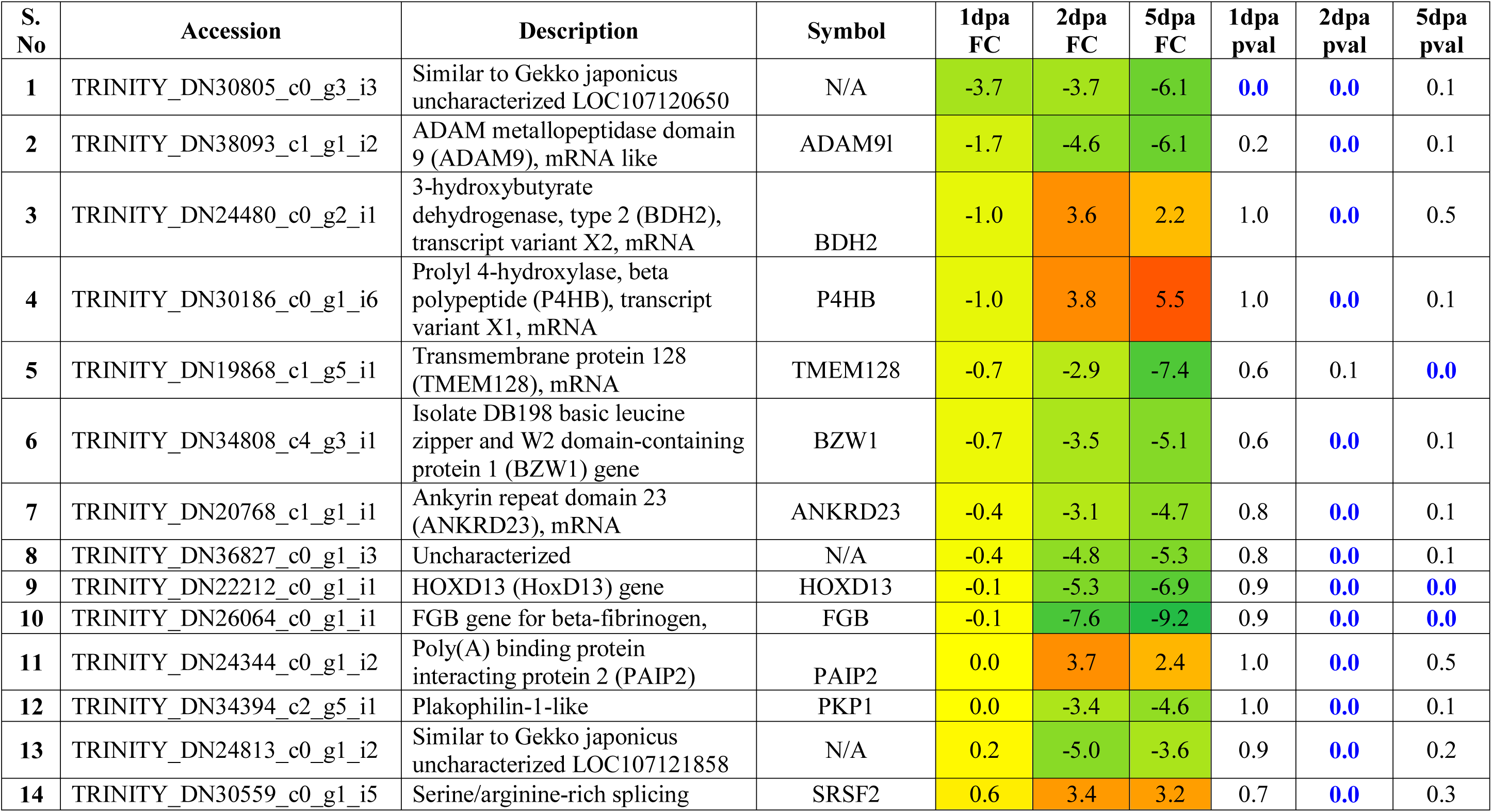
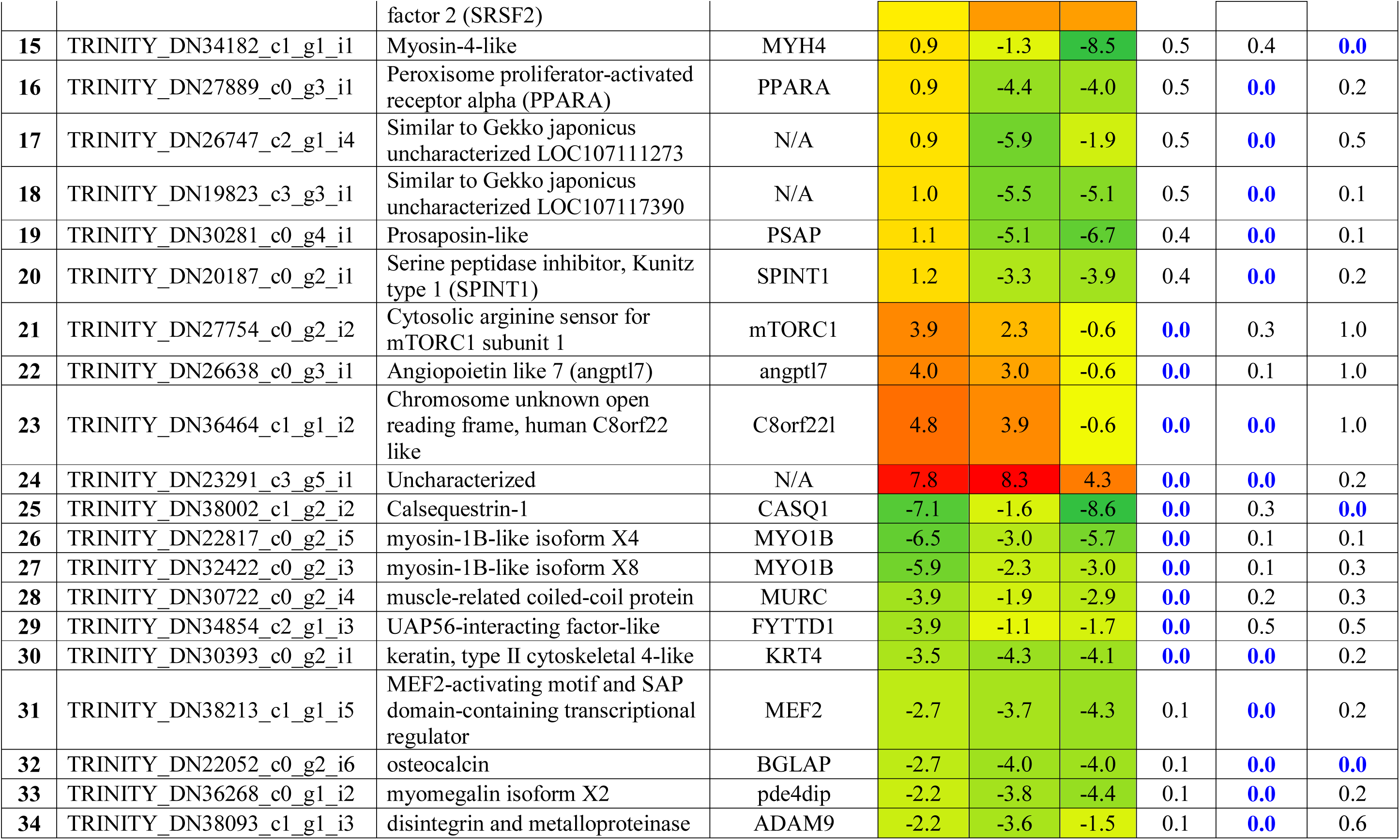
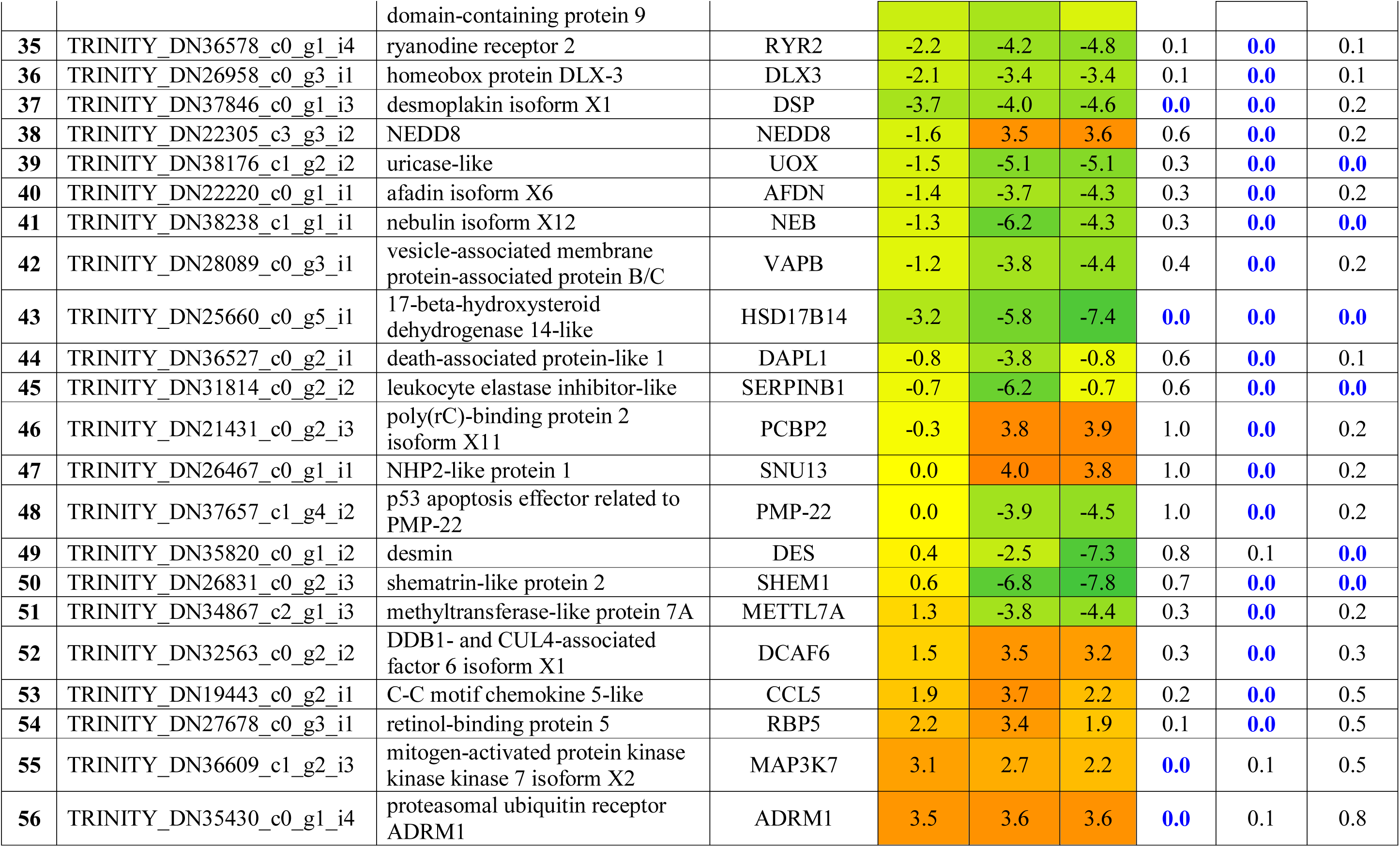
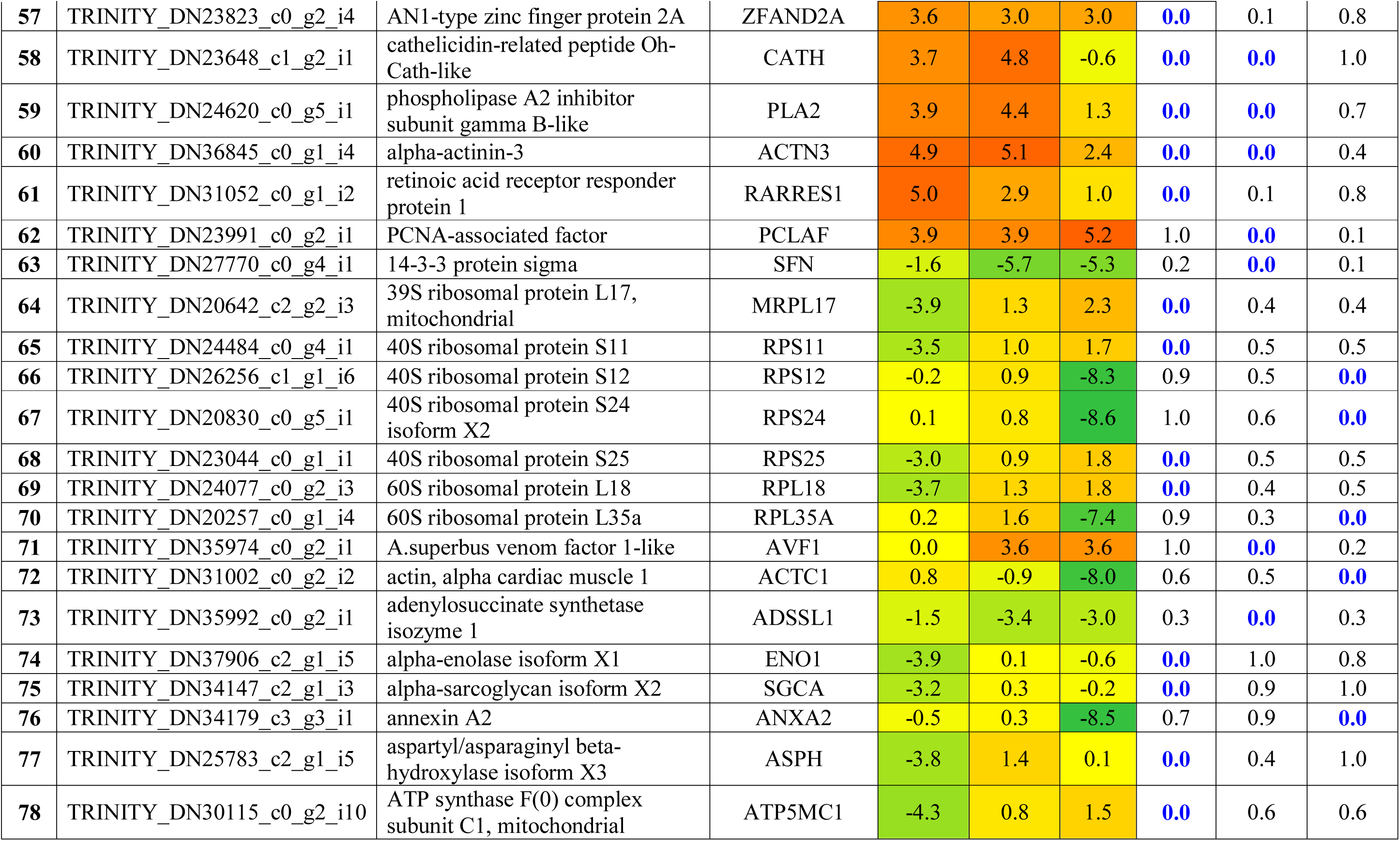
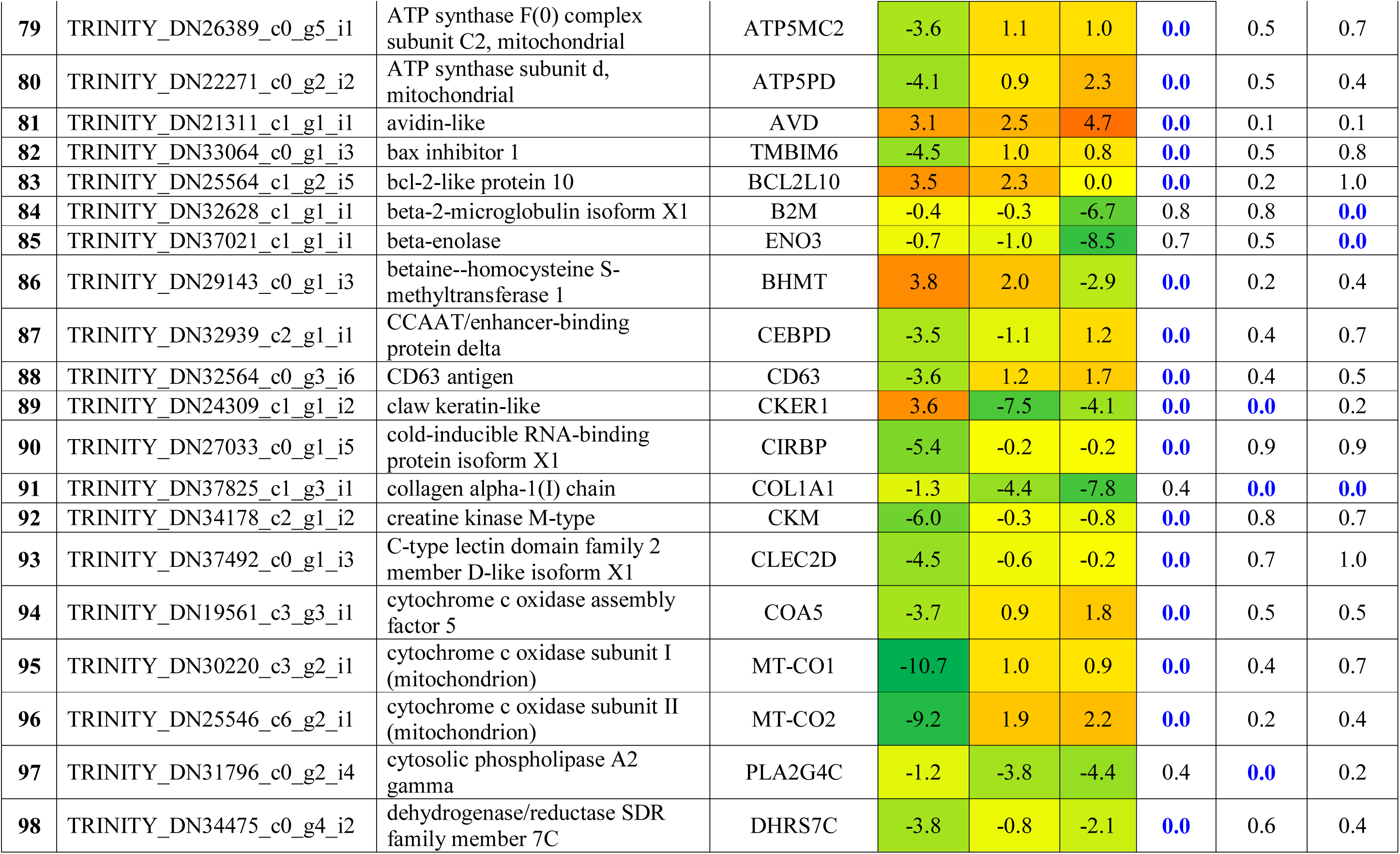
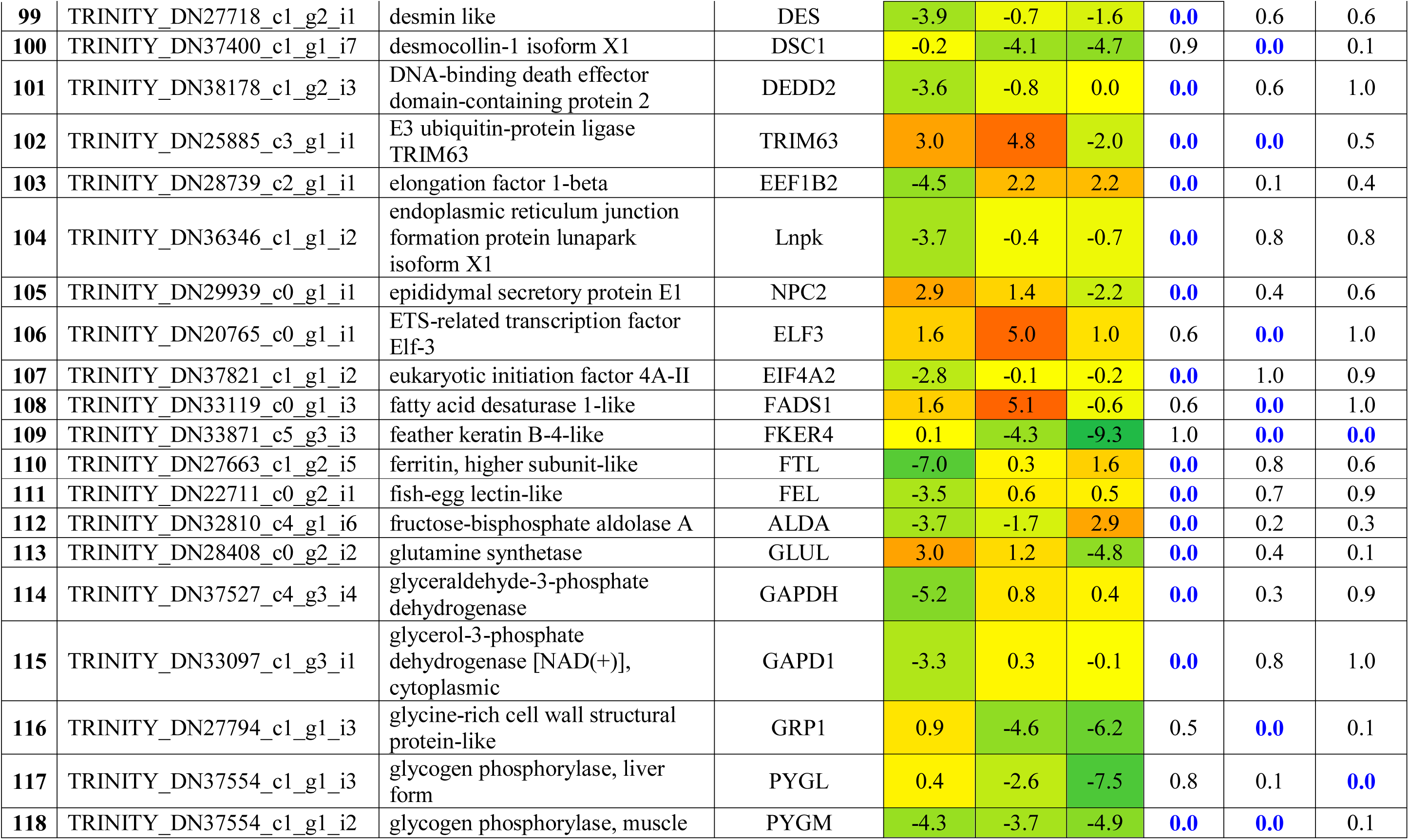
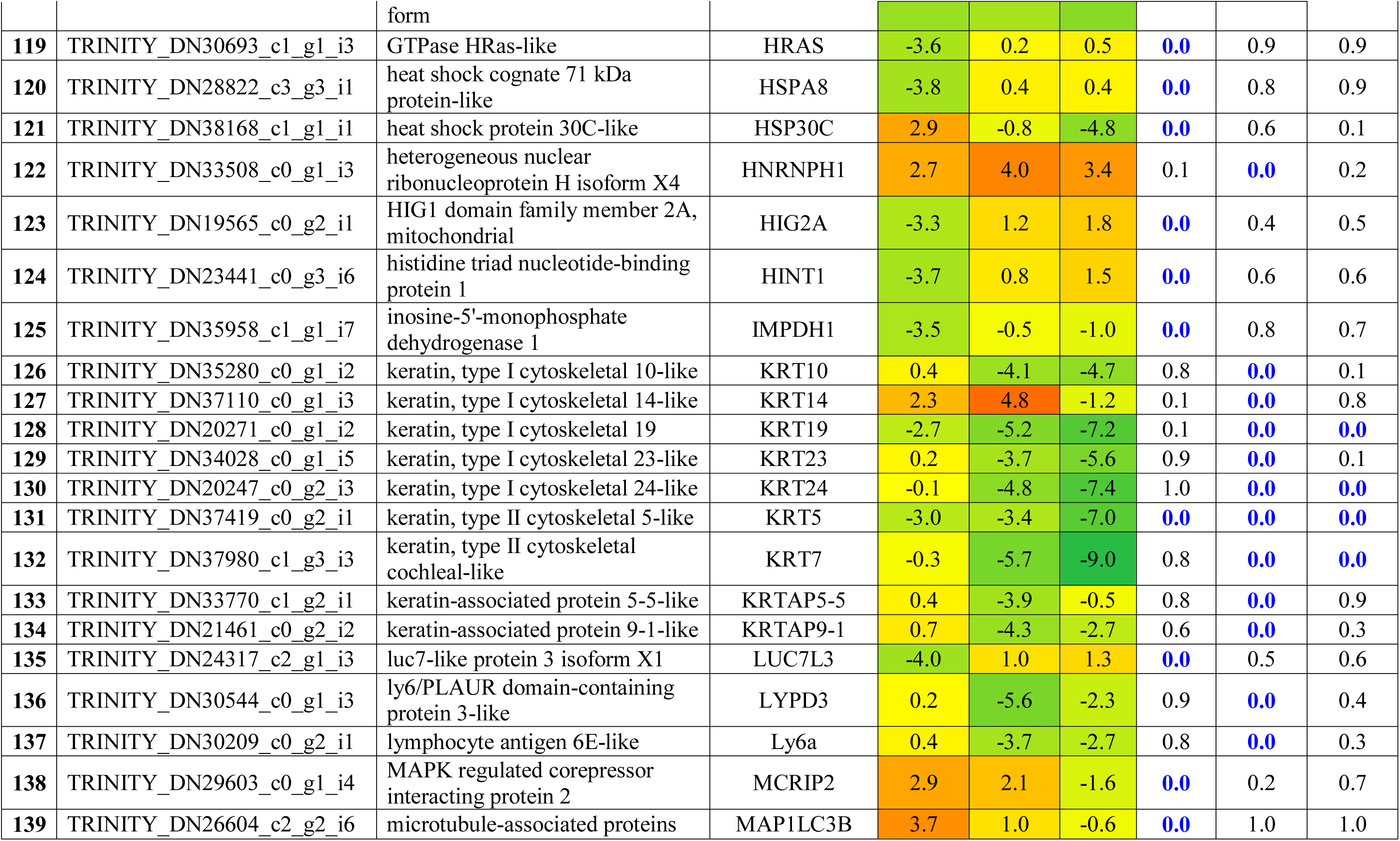
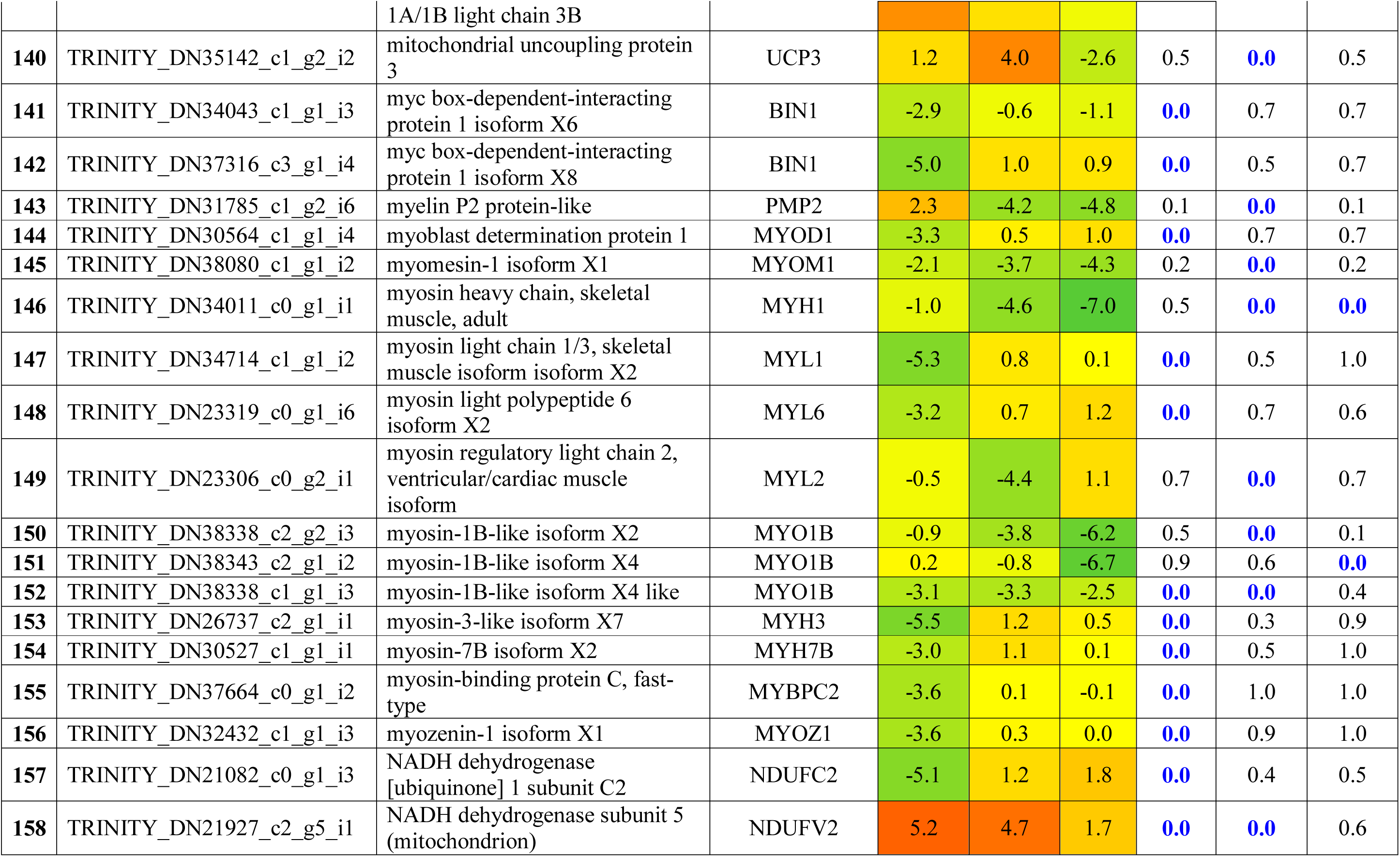
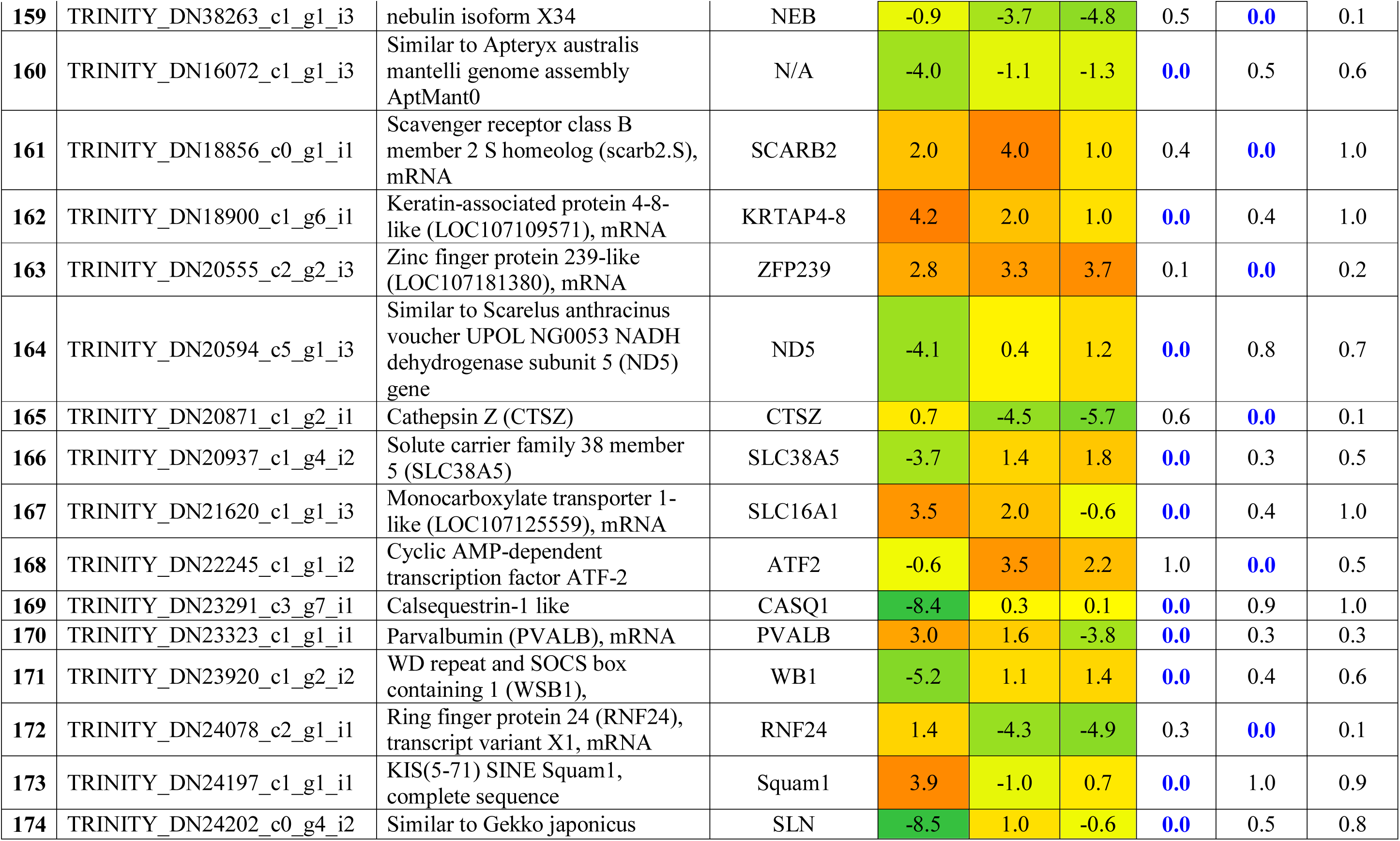
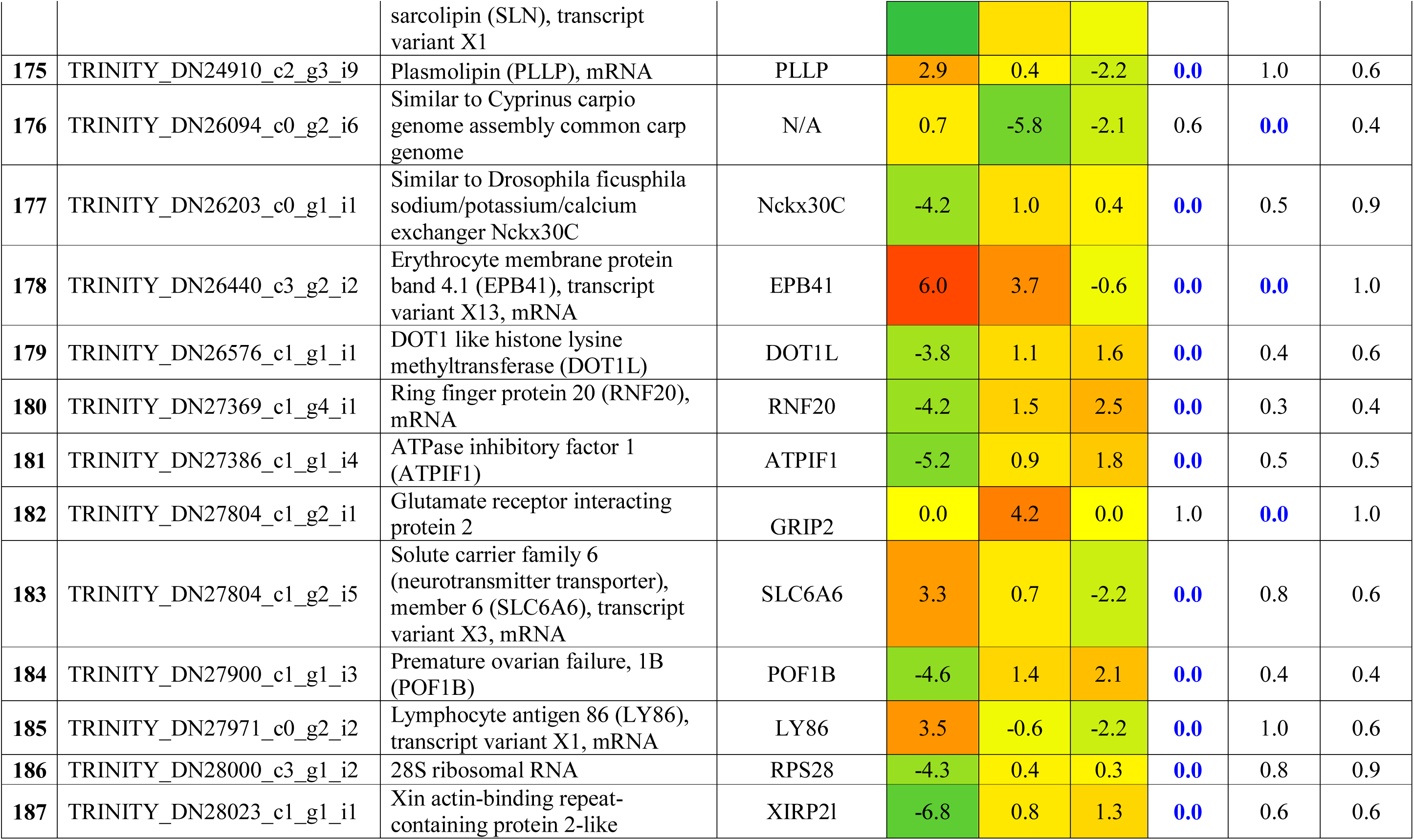
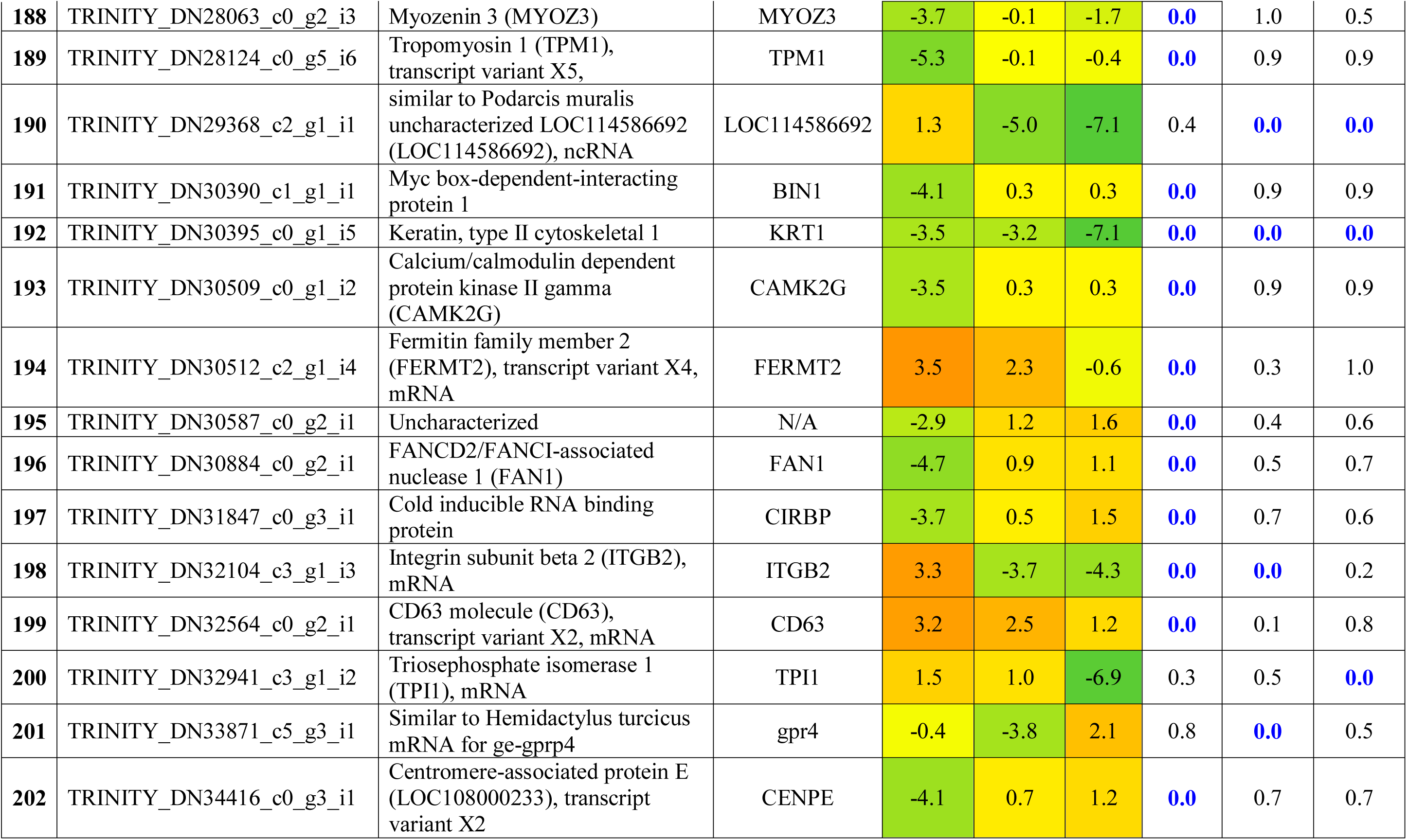
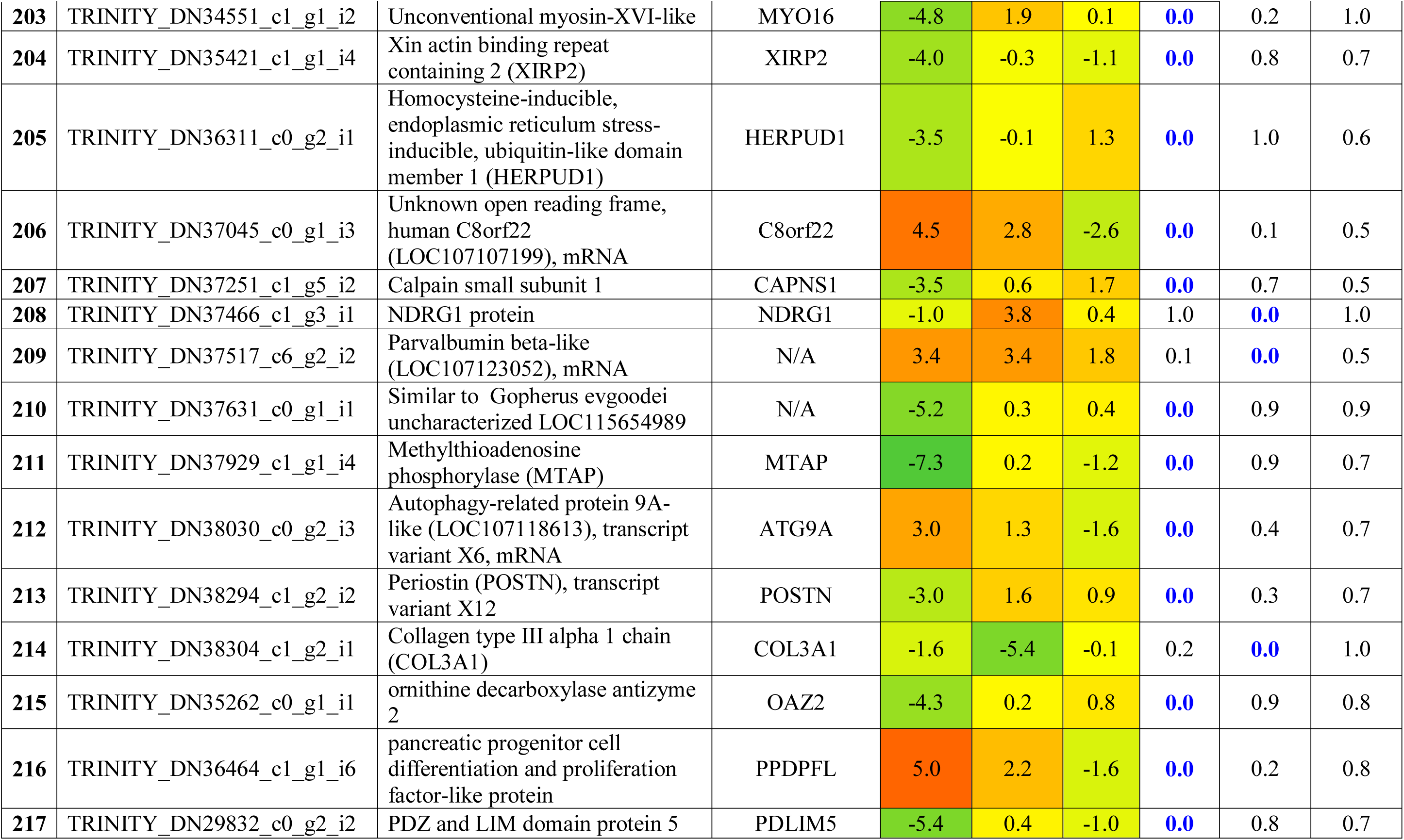
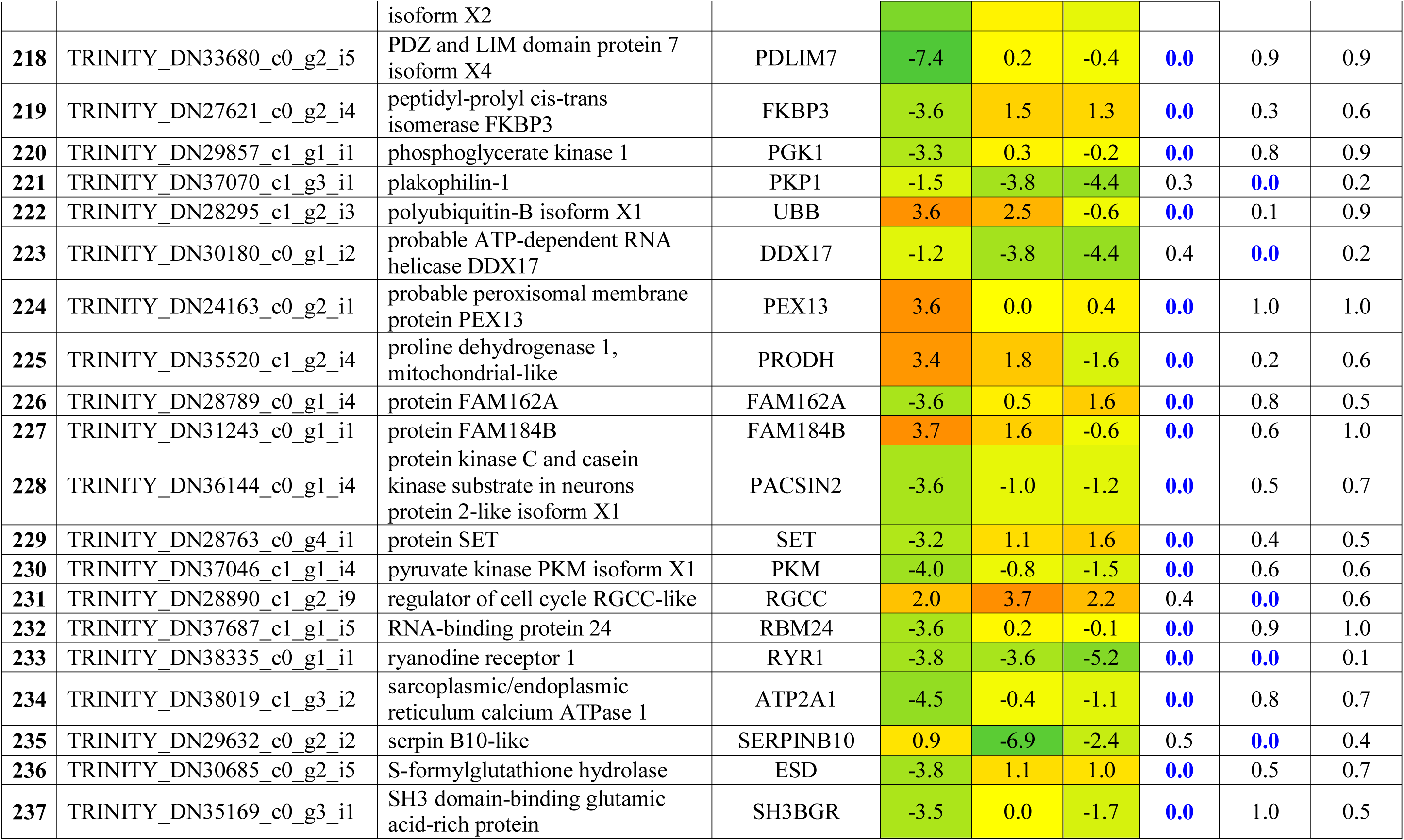
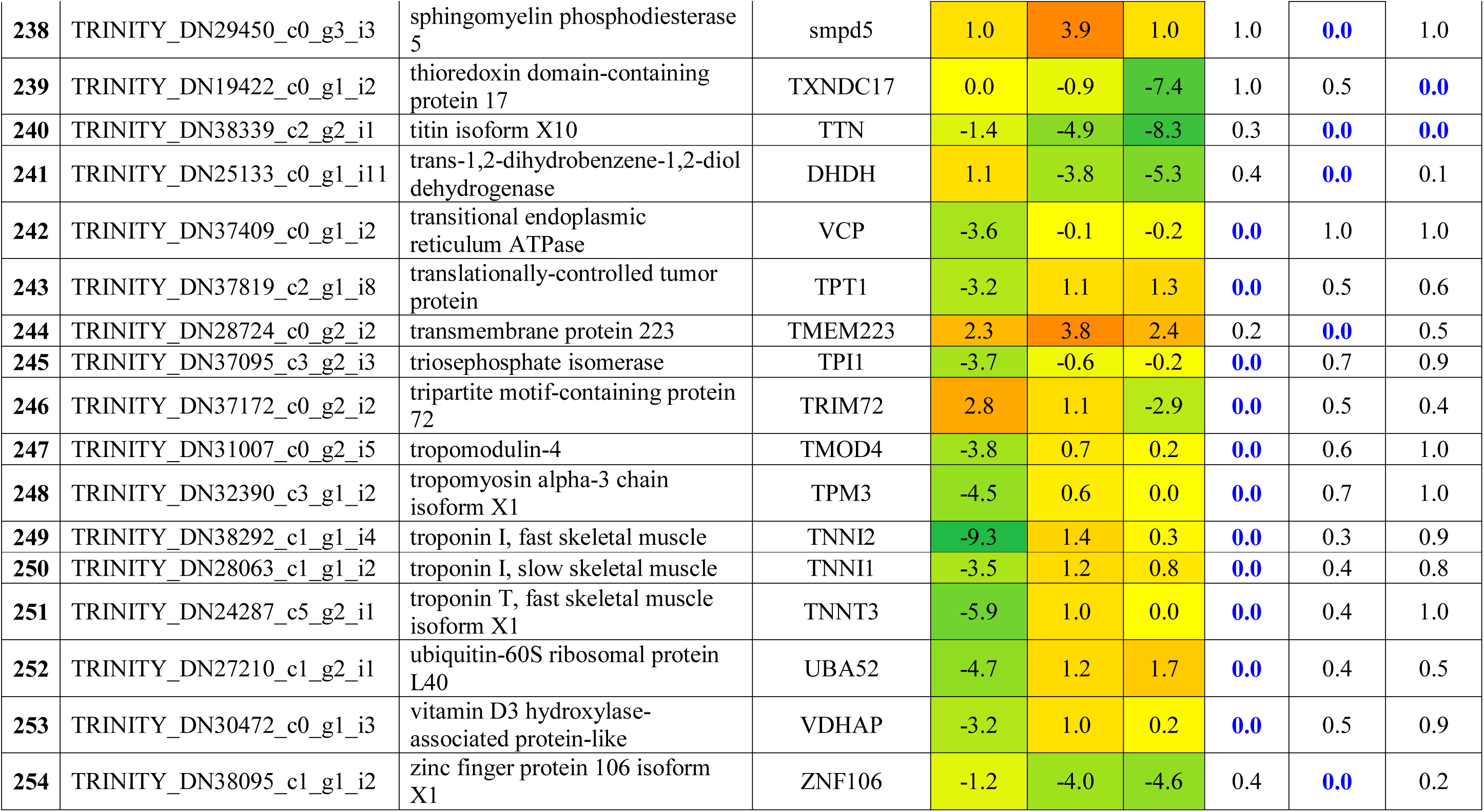
List of differentially expressed genes based on NGS transcriptomic analysis. Accession, description, symbol, 1dpa fold changes, 2dpa fold changes, 5dpa fold changes, 1dpa p value, 2dpa p values and 5dpa p values were shown in each column.

### Proteomic analysis of regeneration

A total of 128 proteins were found to be differentially regulated during the early stage of regeneration of gecko tail tissues for having minimum of one log fold changes significantly (Table 2). Annexin A2, Cathepsin B and MYL1 were some of the major upregulated proteins and Myozenin, GST and GSFK were the major down regulated proteins. A total of 36 proteins which were differentially regulated at the proteome level were also found to be differentially regulated at gene level based on the transcriptomics analysis (Figure 1c). Like transcriptome expression pattern, proteome expressions were also found to be associated with regeneration by clustering of 2 and 5dpa (Figure 1d). All the proteins which were upregulated at 1dpa were found to be down regulated either at 2 or 5 dpa. Similarly all the down regulated proteins were found to be upregulated at 2 or 5dpa.

**Table 2:**
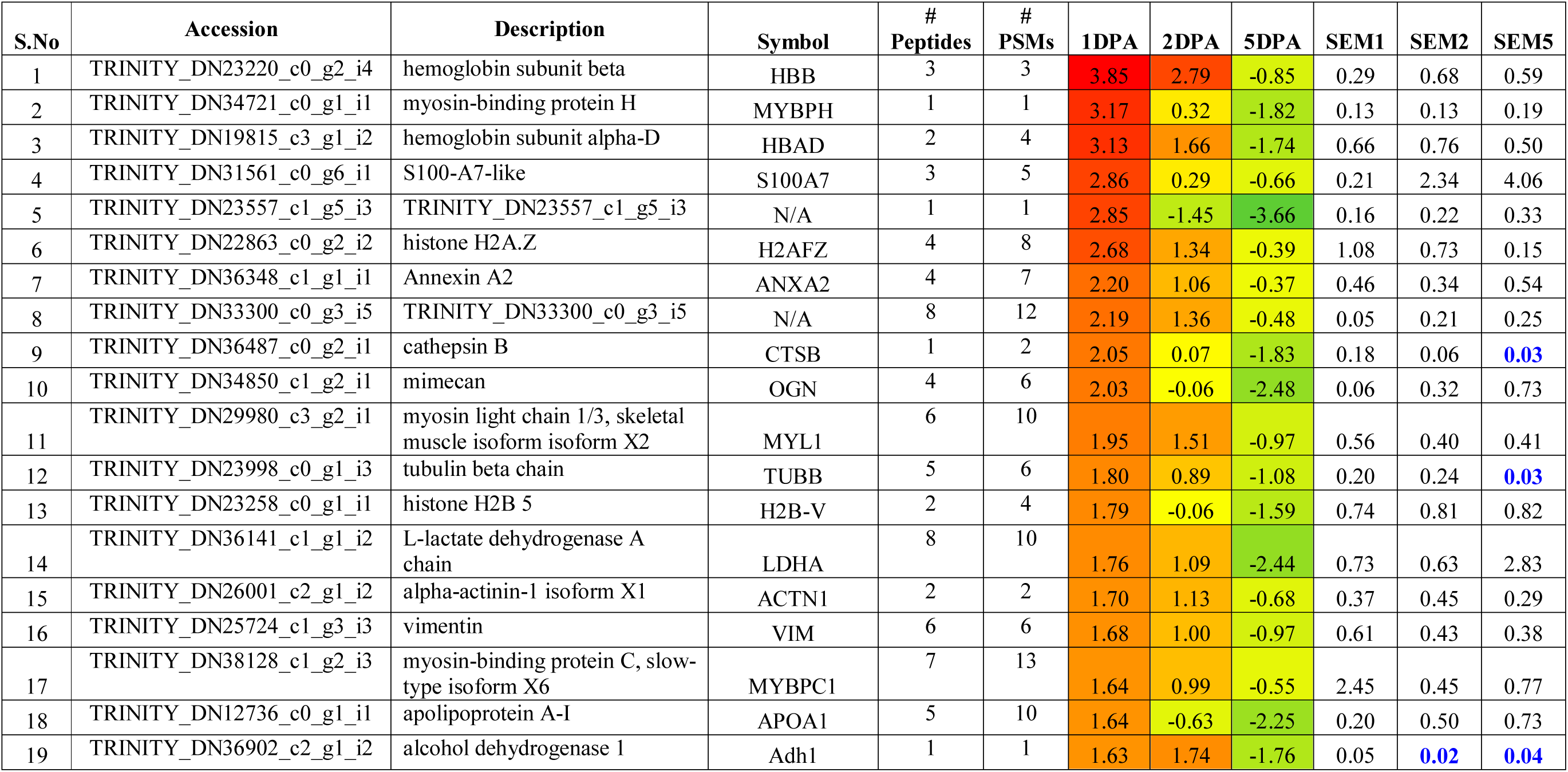
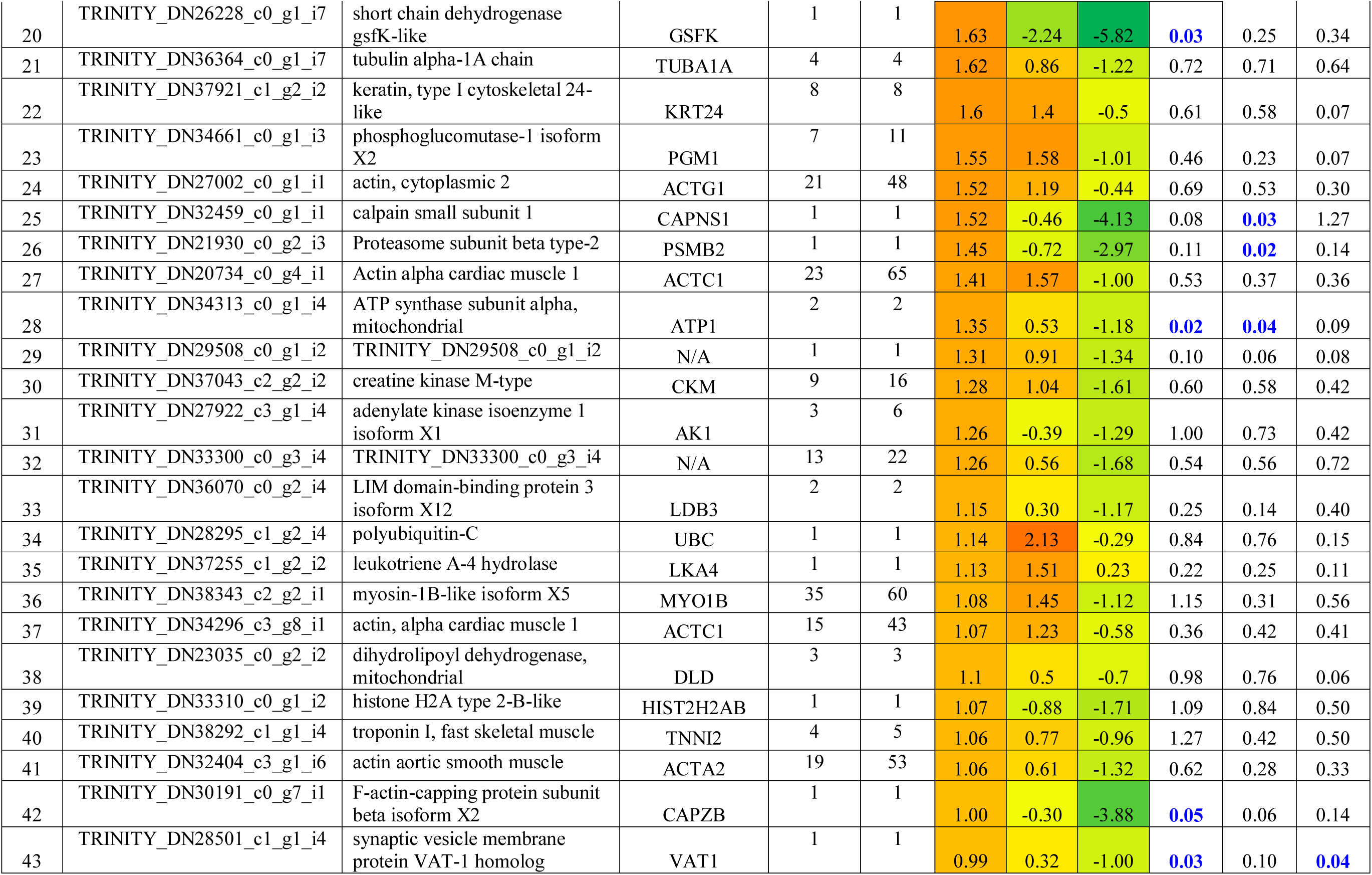
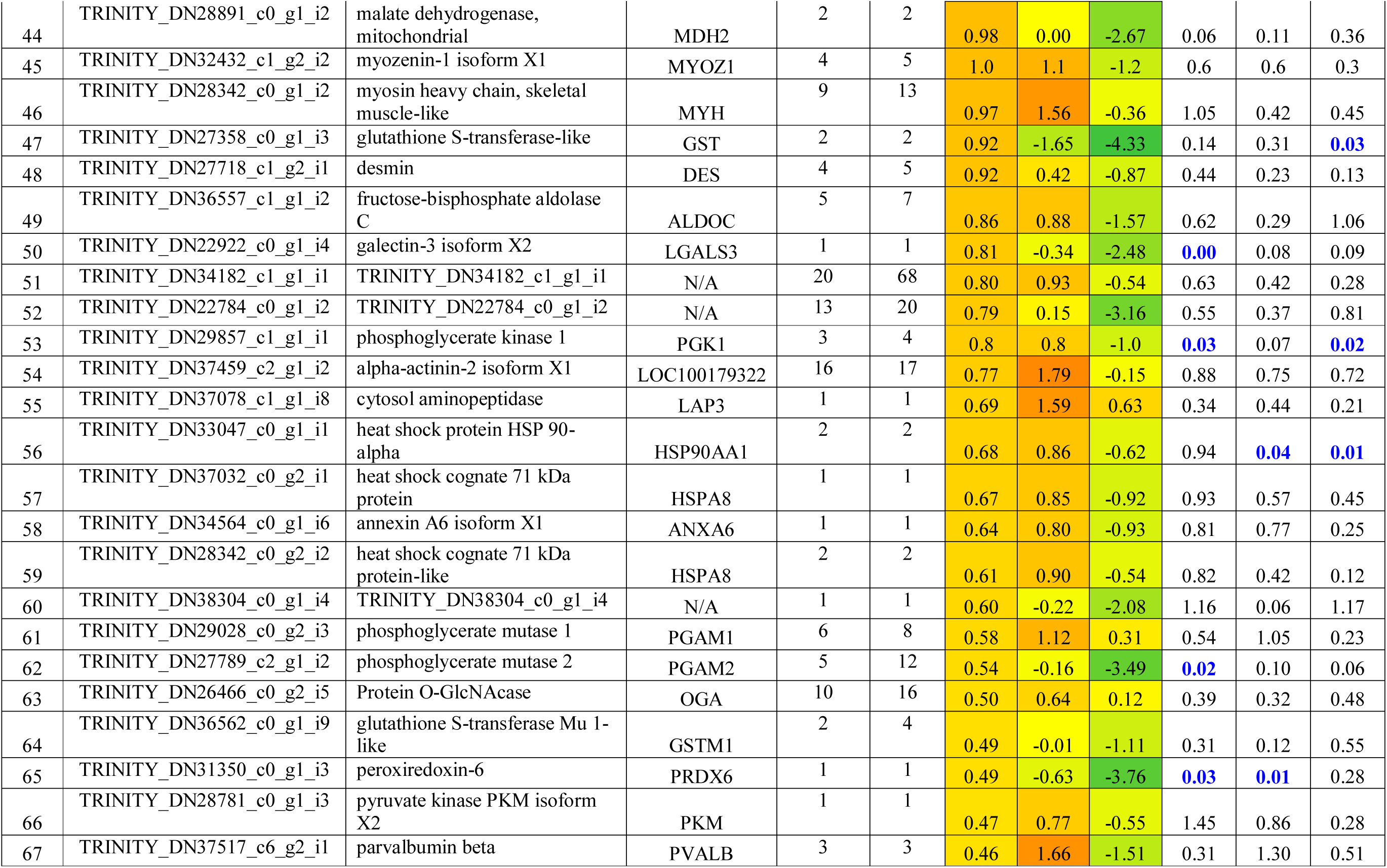
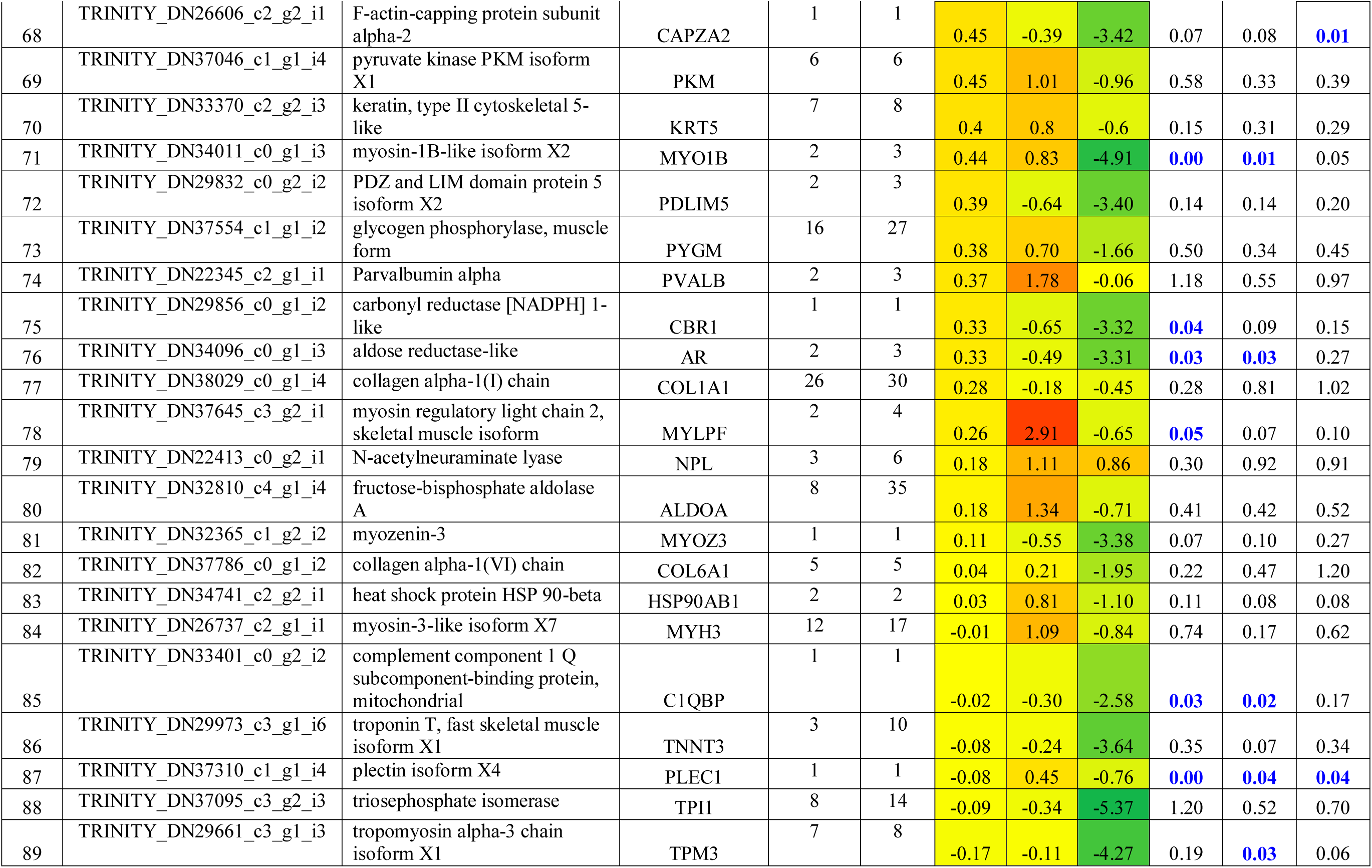
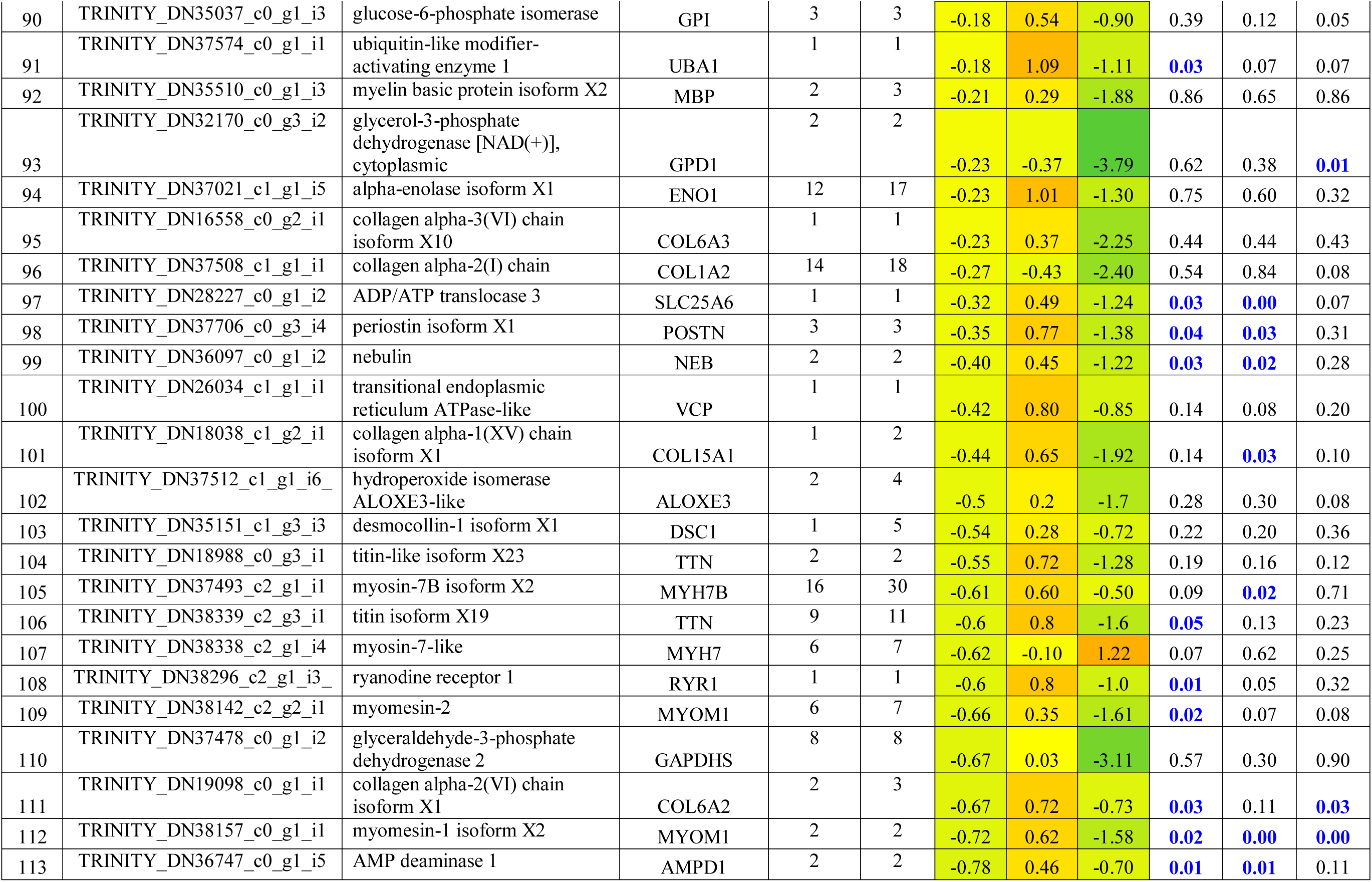
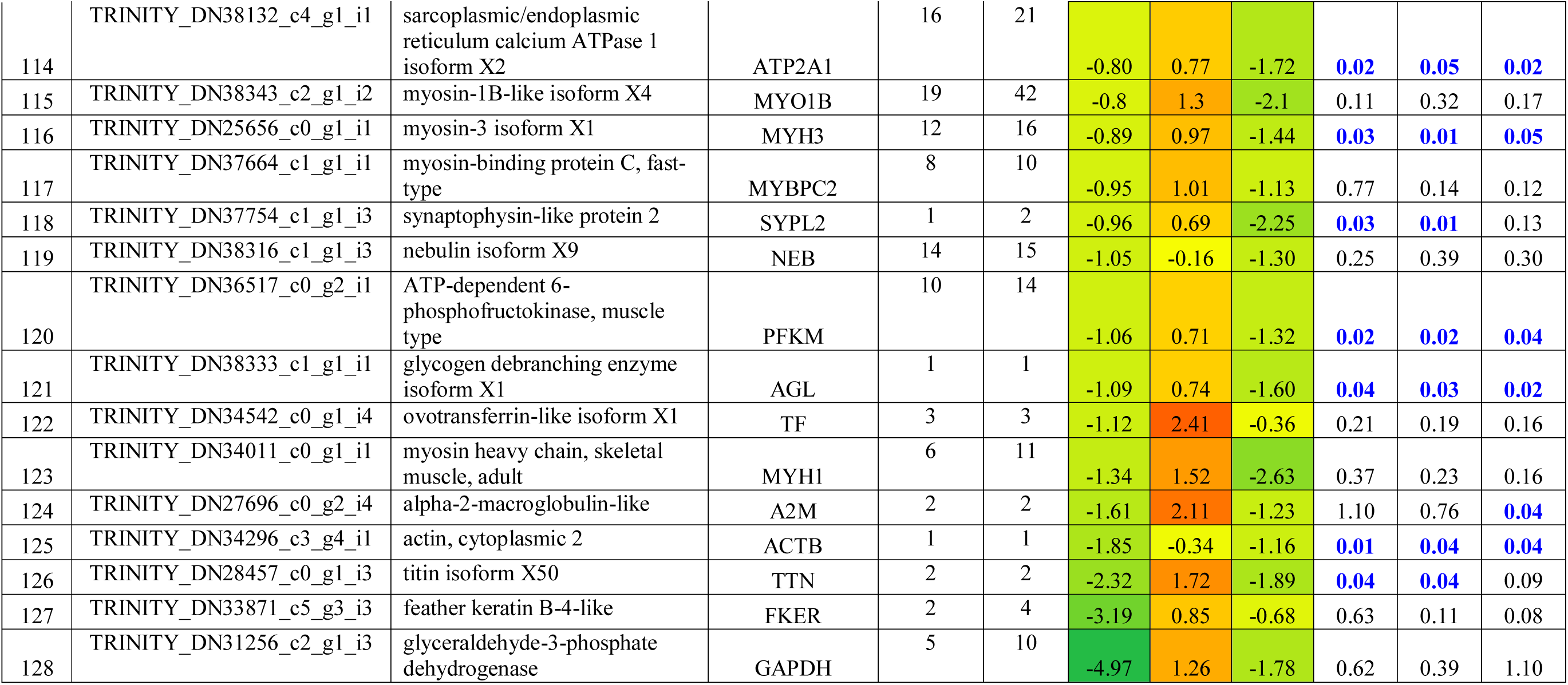
List of differentially expressed protein based on iTRAQ proteomic analysis. Accession, description, symbol, No of peptides identified, peptide sequence matches, 1dpa fold changes, 2dpa fold changes, 5dpa fold changes, 1dpa SEM, 2dpa SEM and 5dpa SEM were shown in each column.

### Validation of gene expression

RTPCR analysis of 50 genes selected for the validation study showed significant differential expression of the genes for the regeneration mechanism (Figure 2a and 2b). Almost all the up and down regulated genes which were identified from the transcriptomic analysis were found to be associated with regeneration through differential regulation based on RTPCR analysis (Figure 1b). Heat map analysis of the gene expression based on RTPCR analysis also showed association of 2 and 5dpa as cluster against 1dpa expression.

**Figure 2:**
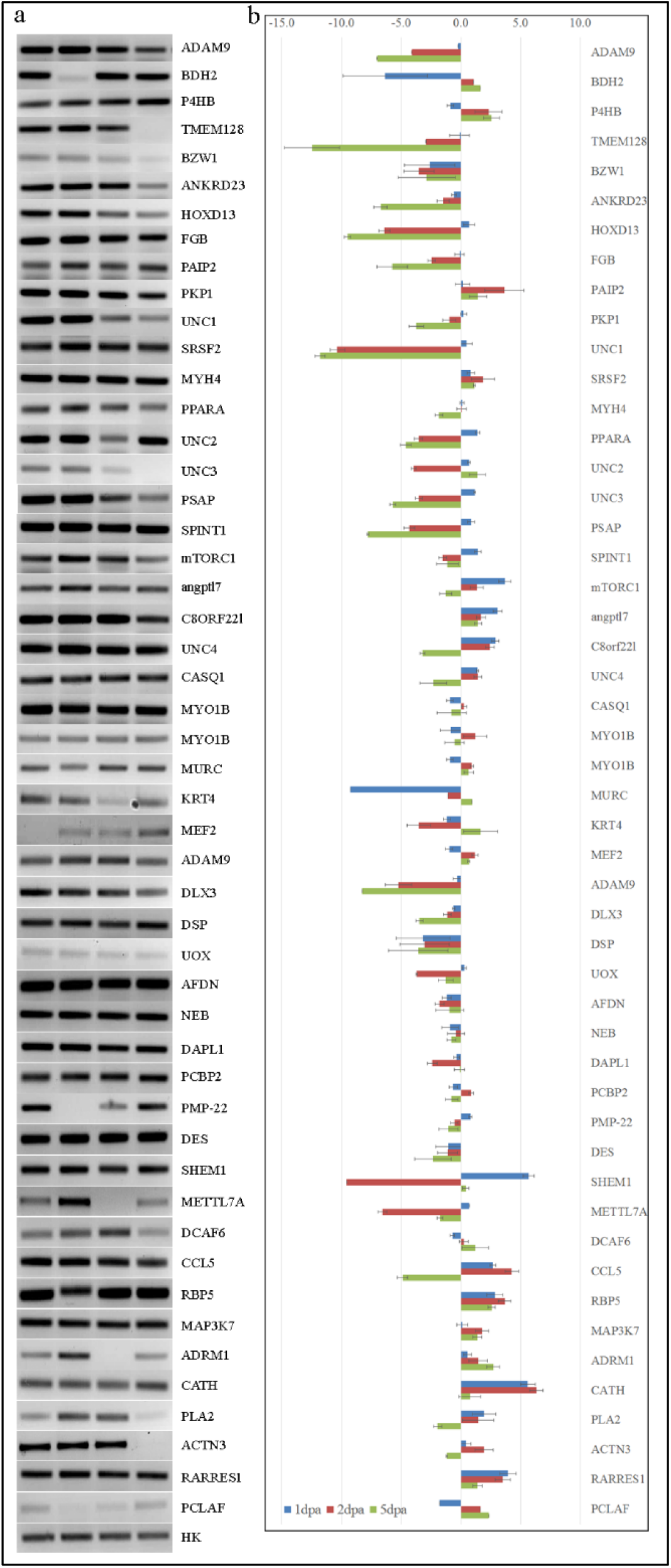
RTPCR analysis of differentially expressed transcripts. a. Gel image of all the 50 genes along with housekeeping gene. b. Bar diagram of expression visualising the up and down regulation of the genes.

### Network Pathway Analysis

A total of 327 genes/proteins were mapped by the Ingenuity Pathway analysis software for the network and pathway analysis. The major molecular and cellular functions associated with the differentially regulated genes/proteins during regeneration are cellular assembly & organization, cellular compromise, cellular functions & maintenance and Cellular development. The major physiological system development and functions associated with regeneration mechanism are skeletal & muscular system development, embryonic development, organ development and tissue development.

The most significant canonical pathways which were found to be associated with genes/proteins were GP6 signaling pathway, Protein kinase A signaling, Telomerase signaling BAG2 signaling, paxiling signaling, VEGF signaling and various metabolic pathways (Figure 3a). The major network pathways associated with the identified and dysregulated genes/proteins based on differential analysis includes Cell morphology and Embryonic development (Figure 3b), Cellular assembly and organization (Figure 3c), Organ and organismal development (Figure 3d) and Skeletal and Muscular development (Figure 3e).

**Figure 3:**
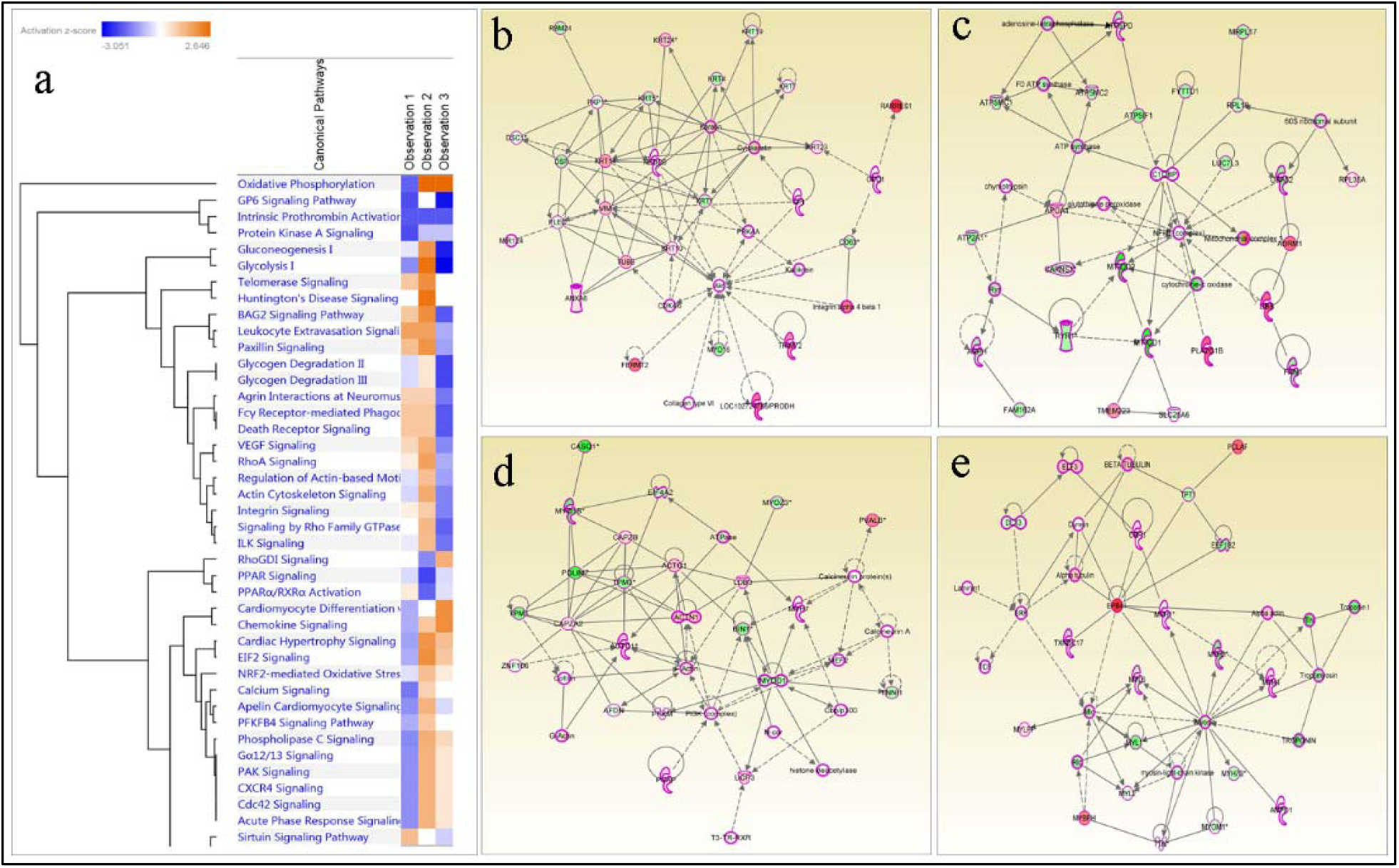
a. Canonical Pathway associated with the differentially expressed genes/proteins for tail regeneration; b. Cell morphology and Embryonic development network pathway; c. Cellular assembly and organization network pathway; d. Organ and organismal development network pathway; e. Skeletal and Muscular development network pathway.

## Discussion

This study aimed to evaluate the genomic and proteomic changes during the early stages of tail tissue regeneration in house lizard. Zebrafish and other invertebrates have been extensively studied for understanding the epimorphic regeneration [27, 28] at the genomic and proteomic level. This study has been done similar to our earlier study using the house gecko as the model animal.

The study not only mapped the transcriptome of gecko tail tissue but also identified the list of genes and proteins involved in regeneration of the tail tissue upon amputation. Several known and unknown genes/proteins which were identified in this study were also involved in epimorphic regeneration of caudal fin tissue of zebrafish [27]; arm of brittle star [28] and tail of Hemidactylus flaviviridis [12]. 417 genes and 134 proteins were identified as differentially regulated in the regenerating tissues. Validation of 50 genes identified from the transcriptomic analysis confirms the differential regulation of the genes for the biomechanism of regeneration, which is also evident from the hierarchical heat map analysis of the transcriptome and proteome and RTPCR expression analysis.

The canonical pathway associated with the differentially expressed genes/proteins are mostly signalling and metabolic pathways. The major network pathways associated with the tail tissue regeneration are Cell morphology, embryonic development and skin development (Figure 3b) involving 26 genes/proteins of the study such as DSC1, DSC2, GPI, DSP and keratin proteins. Similarly, 25 genes/proteins, such as ADRM1, APOA1, ASPH, ATP2A1 were associated with Cellular assembly and organization (Figure 3c). Organ and organismal development network pathway (Figure 3d) were associated with 28 genes/proteins such as ACTC1, ACTG1, BIN1, CAPZA2 and CAPZB. Skeletal and muscular system development network pathway (Figure 3e) was associated with 25 genes/proteins such asCBR1, DLX3, EEF1B2 and DLX3. This study has identified and associated various genes/proteins and their network pathways for the tail tissue regeneration of lizard which were also found to be associated with epimorphic regeneration of organs in other animals as in zebrafish [27] and echinoderms [28]. Further understanding of the differential expressed genes/proteins and their regulation might lead to better insight in understanding the regenation of the gecko tail.

## Acknowledgements

This work was supported by CSIR-YSA Project. The authors are grateful to Noorul Fowzia for critically reviewing the manuscript.

